# Alternatively-Spliced CAMKK2 isoforms drive differential metabolic stress response and the regulation of ferroptosis

**DOI:** 10.1101/2025.05.13.653837

**Authors:** Sze Cheng, Hyunjoo Kim, Naima Ahmed Fahmi, Meeyeon Park, Marella Canny, Ritika Kukreja, Ethan J. Beckermann, Michael Latham, Wei Zhang, Jeongsik Yong

## Abstract

Calcium/Calmodulin-dependent protein kinase kinase 2 (CAMKK2) is a multifunctional kinase that regulates metabolic processes by phosphorylating downstream targets. CAMKK2 is expressed as several highly similar protein isoforms, although the specific functions of these individual isoforms remain largely unexplored. These isoforms have been shown to display tissue-specific expression patterns, suggesting unique roles in distinct cellular contexts. In this study, we investigated the biochemical and functional relevance of CAMKK2 isoforms, LF and SF, in the context of glucose metabolism. Co-immunoprecipitation experiments revealed that the LF isoform preferentially binds the adaptor protein 14-3-3, while the SF isoform interacts with Calmodulin. Furthermore, the interaction between SF and Calmodulin was enhanced upon glucose starvation, whereas this interaction was not observed for LF. To assess their functional significance, we generated doxycycline-inducible, isoform-specific HeLa cell lines. Under low glucose conditions, cells expressing the LF isoform failed to activate AMPK, while cells expressing the SF isoform exhibited robust increase in AMPK phosphorylation. Moreover, LF-expressing cells accumulated higher levels of reactive oxygen species, upregulated ferroptosis-related genes via BACH1/NRF2, and displayed increased cell death compared to SF-expressing cells. Collectively, these findings demonstrate that CAMKK2 isoforms exhibit differential selectivity for protein partners and mediate distinct downstream signaling pathways, impacting metabolism in cancer.

## Introduction

Upon transcription, nascent pre-mRNAs undergo alternative splicing, a process by which different combinations of exons are selectively included or excluded to generate multiple mRNA isoforms^1,2^. This mechanism is prevalent in over 90% of human genes and enables eukaryotic cells to produce a diverse array of protein isoforms from a single gene^3,4^. This RNA processing enriches the proteome with functional variety and increased complexity. Alternative splicing is meticulously regulated by the interactions between trans-acting splicing factors and cis-acting elements on the pre-mRNA^5,6^. This control is both tissue-specific and context-dependent, rendering the splicing landscape highly dynamic^5,6^. Such precise modulation allows cells to adapt their transcript isoform profiles to various physiological conditions^7,8^. Changes in alternative splicing have been linked to disease progression^9–11^, and recent research has elucidated how it is influenced by the cellular mTOR signaling pathway^12–15^.

The mTOR signaling pathway acts as a pivotal metabolic controller, responding to a variety of extracellular signals to manage numerous cellular functions, including protein synthesis, cell growth, and metabolism^16,17^. In cancer, mTOR signaling is frequently upregulated to support the increased demands for biomass production^18,19^. Recent studies have not only highlighted the impact of mTOR on alternative splicing throughout the transcriptome but have also demonstrated its role in modifying pre-mRNA polyadenylation. For instance, mTOR activation can promote the production of 3’-untranslated region (UTR) shortened transcripts by selecting proximal polyadenylation signals in the 3’-UTR, thereby enhancing protein synthesis^13^. Additionally, our latest research indicates that changes in mTOR signaling in cells correlate with a dichotomous expression profile of intron polyadenylated transcripts, a pattern also seen in cancer patient transcriptomes^15^.

Beyond its conventional role in protein synthesis and cellular anabolism, our findings establish a novel link between mTOR signaling and proteome diversification through transcriptome alterations (i.e. transcriptome plasticity)^14^. Although the relationship between mTOR signaling and proteome diversification is evident, the specific functional impacts of these changes remain to be fully elucidated.

Calcium/Calmodulin-dependent protein kinase kinase 2 (CAMKK2) is a calcium-sensitive kinase that regulates energy homeostasis, inflammation, and gene regulation by phosphorylating a diverse set of substrates, such as AMPK and CAMKIV^20^. Its kinase activity has been shown to be influenced by its binding to calcium-bound Calmodulin (CaM)^20^. Dysregulation of CAMKK2 driven by overexpression or genetic mutations is implicated in diabetes, neurodegeneration, and cancer^21–25^. CAMKK2 exists as several highly similar protein isoforms generated by alternative splicing and polyadenylation^26^. Due to the high sequence homology and domain conservation, CAMKK2 isoforms are largely treated as the same protein. However, one notable study revealed isoform-specific functions in neuronal differentiation^26^, suggesting that CAMKK2 alternative splicing isoforms have distinct roles. Additionally, previous studies have shown that cancer cells exclusively express one CAMKK2 isoform compared to normal, healthy tissues^27,28^. While largely unexplored, these previous findings suggest the specialized roles of CAMKK2 isoforms in distinct cellular context.

In this study, we explore how mTOR-regulated protein isoform production impacts cellular processes. Specifically, we demonstrate that mTOR-regulated alternative splicing of CAMKK2 leads to paradoxical functions of this gene in response to metabolic stress, highlighting a complex scenario where a single gene can exhibit contrasting roles depending on the context.

## Results

### Cellular context-dependent alternative splicing *of CAMKK2* isoforms

Our previous research demonstrated that cellular context regulated by mTOR signaling modulates transcriptome-wide alternative splicing^14^. Specifically, we identified that elevated mTOR activity induces widespread exon skipping events. A significant example is observed in mTOR-hyperactivated mouse embryonic fibroblasts (*Tsc1*^−/-^ MEFs), where *Camkk2* undergoes alternative splicing that results in a transcript isoform lacking exon 16 (Fig.1a). This exon skipping event is notably sensitive to cellular mTOR signaling, evidenced by significantly reduced exon skipping when mTOR activity is pharmacologically inhibited with Torin 1, or when evaluated in corresponding wild-type (WT) MEFs (Fig. 1a and 1b).

**Figure 1.**
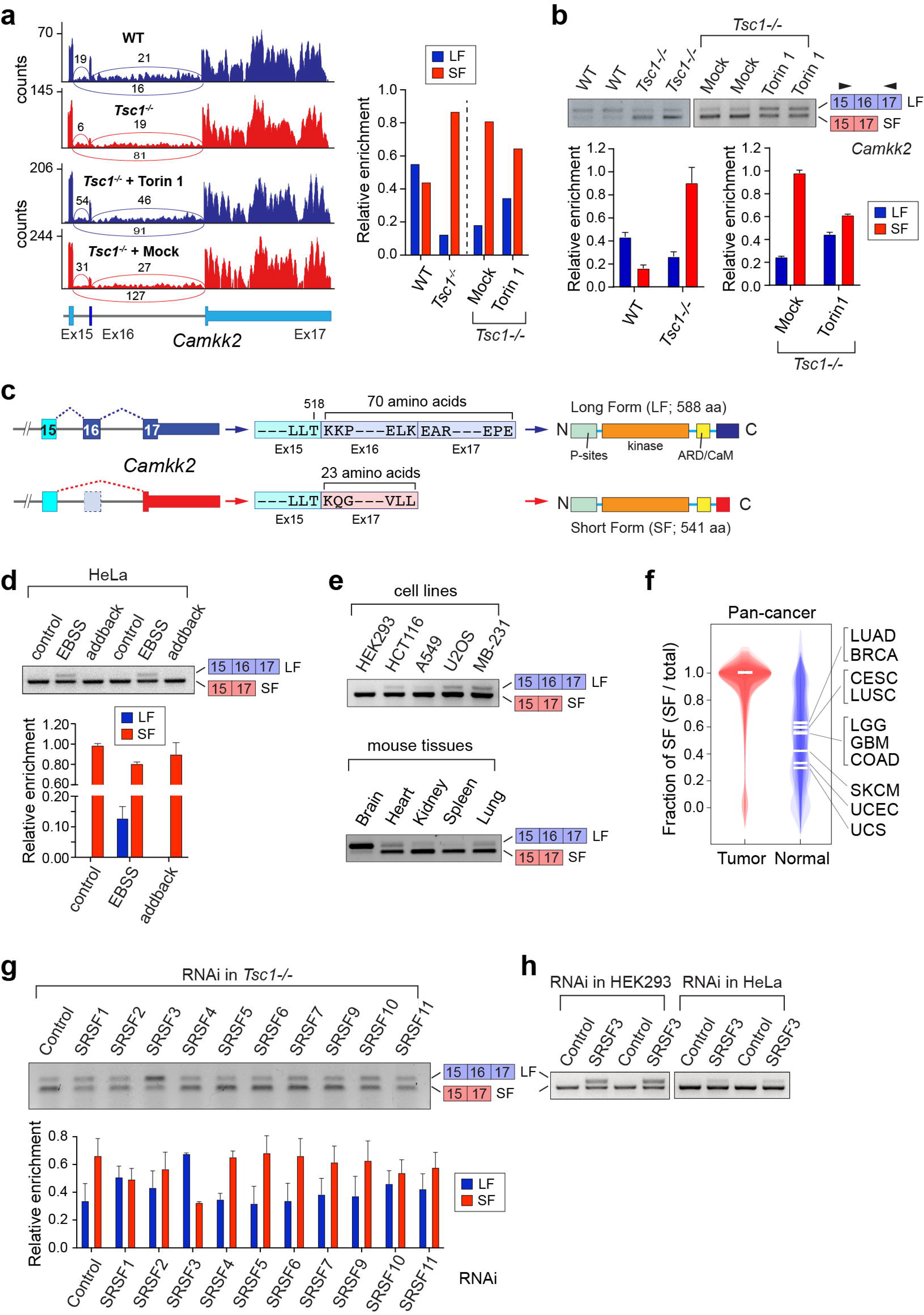
Cellular context-dependent alternative splicing (AS) *of CAMKK2* isoforms. **a**) RNA-seq read alignments of *Camkk2* exon 15 to 17 region in wild-type (WT), *Tsc1^−/-^*, mock-treated, and Torin 1-treated *Tsc1^−/-^* mouse embryonic fibroblasts (MEFs). The relative enrichment of the isoforms was calculated using the read coverage from RNA-seq. **b**) Semi-quantitative RT-PCR analyses of *Camkk2* exon 16 AS in wild-type (WT), *Tsc1^−/-^*, mock-treated, and Torin 1-treated *Tsc1^−/-^* MEFs. The same analysis was performed in HeLa cells cultured in a nutrient-rich (control and add-back samples) and -deprived environment (EBSS sample) **d**) and various human and mouse immortalized cell lines and tissues (**e**). **c**) Schematic depiction of CAMKK2 exon 16-included and -excluded isoform protein sequences. **f**) A super-imposed violin plot of the CAMKK2 expression profile in The Cancer Genome Atlas (TCGA) and The Genotype-Tissue Expression (GTEx) project. The fraction of short isoform (SF) over total *CAMKK2* expression is shown. LUAD; Lung adenocarcinoma, BRCA; Breast invasive carcinoma, CESC; Cervical squamous cell carcinoma and endocervical adenocarcinoma, LUSC; Lung squamous cell carcinoma, LGG; Low-grade glioma, GBM; Glioblastoma, COAD; Colon adenocarcinoma, SKCM; Skin cutaneous melanoma, UCEC; Uterine corpus endometrial carcinoma, UCS; Uterine carcinosarcoma. p<1.78e-36. **g**) Semi-quantitative RT-PCR analyses of *Camkk2* AS in an RNAi knockdown of SR proteins in *Tsc1^−/-^* MEFs. (n=3). **h**) Semi-quantitative RT-PCR analyses of *CAMKK2* AS in control and SRSF3 knockdown in HEK293 and HeLa cells.

This alternative splicing event, conserved between mouse and human *CAMKK2* genes, produces two distinct isoforms differing only in their C-terminal regions due to a frameshift resulting from exon exclusion (Fig. 1c). Although previous studies have noted the predominant expression of the shorter CAMKK2 isoform in cancer cells^26,27^, many investigations have not rigorously distinguished between these isoforms. Given the substantial sequence homology and shared functional domains between the isoforms, subtle yet potentially significant differences have largely been overlooked in prior research. Our initial investigation across diverse biological contexts revealed selective upregulation of the longer CAMKK2 isoform under serum deprivation conditions in HeLa cervical cancer cells, correlating with mTOR-regulated alternative splicing events (Fig. 1d). Furthermore, variations in CAMKK2 isoform ratios were observed among different cell lines and tissues, with a tendency towards shorter isoform expression in cancer cells and various tissues; notably, the brain predominantly expresses the longer isoform (Fig. 1e).

Further analysis using RNA-seq data from The Cancer Genome Atlas (TCGA) indicated that tumor samples from 26 different cancer types predominantly expressed the short isoform (Fig. 1f). These observations imply that CAMKK2 isoforms expression is context-dependent, tissue-specific, and potentially relevant to cancer progression.

To elucidate mechanisms governing *Camkk2* splicing, we assessed the impact of SR-rich splicing factors in Tsc1-/-MEFs. Knockdown experiments revealed that reducing SRSF3 expression significantly enhanced exon 16 inclusion, whereas knockdown of other SR proteins did not significantly influence *Camkk2* alternative splicing (Fig. 1g). This suggests that SRSF3 specifically suppresses exon 16 inclusion in hyperactivated mTOR contexts. Consistent results were observed in human HEK293 and HeLa cells, reinforcing the evolutionarily conserved suppressive role of SRSF3 on *CAMKK2* exon 16 inclusion (Fig. 1h). Notably, our prior research found SRSF3 upregulation through mTOR-mediated 3’-UTR alternative polyadenylation^13^. Collectively, our findings highlight a conserved regulatory axis involving mTOR signaling and SRSF3-mediated alternative splicing of *CAMKK2*, favoring the expression of shorter CAMKK2 isoform in cancer cells.

### Impact of CAMKK2 isoform-specific C-terminal sequences on protein interactions

CAMKK2 is known to interact with Ca^2+^-Calmodulin (CaM)^20,29,30^, and the critical phosphorylation sites that regulate these interactions have been previously identified^29,31,32^. However, the influence of the distinct C-terminal sequences that differ among CAMKK2 isoforms on their protein interactions has been largely unexplored. To investigate whether these isoform-specific C-terminal sequences influence affect CaM binding, we conducted co-immunoprecipitation experiments to compare the binding of CaM to the long (LF) and short (SF) CAMKK2 isoforms. Flag-tagged CAMKK2 isoforms were transiently expressed, immunoprecipitated in the presence of Ca^2+^, and their interactions with endogenous CaM were analyzed via western blotting. Notably, we observed a significantly enhanced CaM binding to CAMKK2-SF than to CAMKK2-LF (Fig. 2a). Given that both isoforms share an identical CaM-binding domain, this stronger binding suggests that the unique C-terminal region of CAMKK2-SF plays a critical role in facilitating CaM interaction.

**Figure 2.**
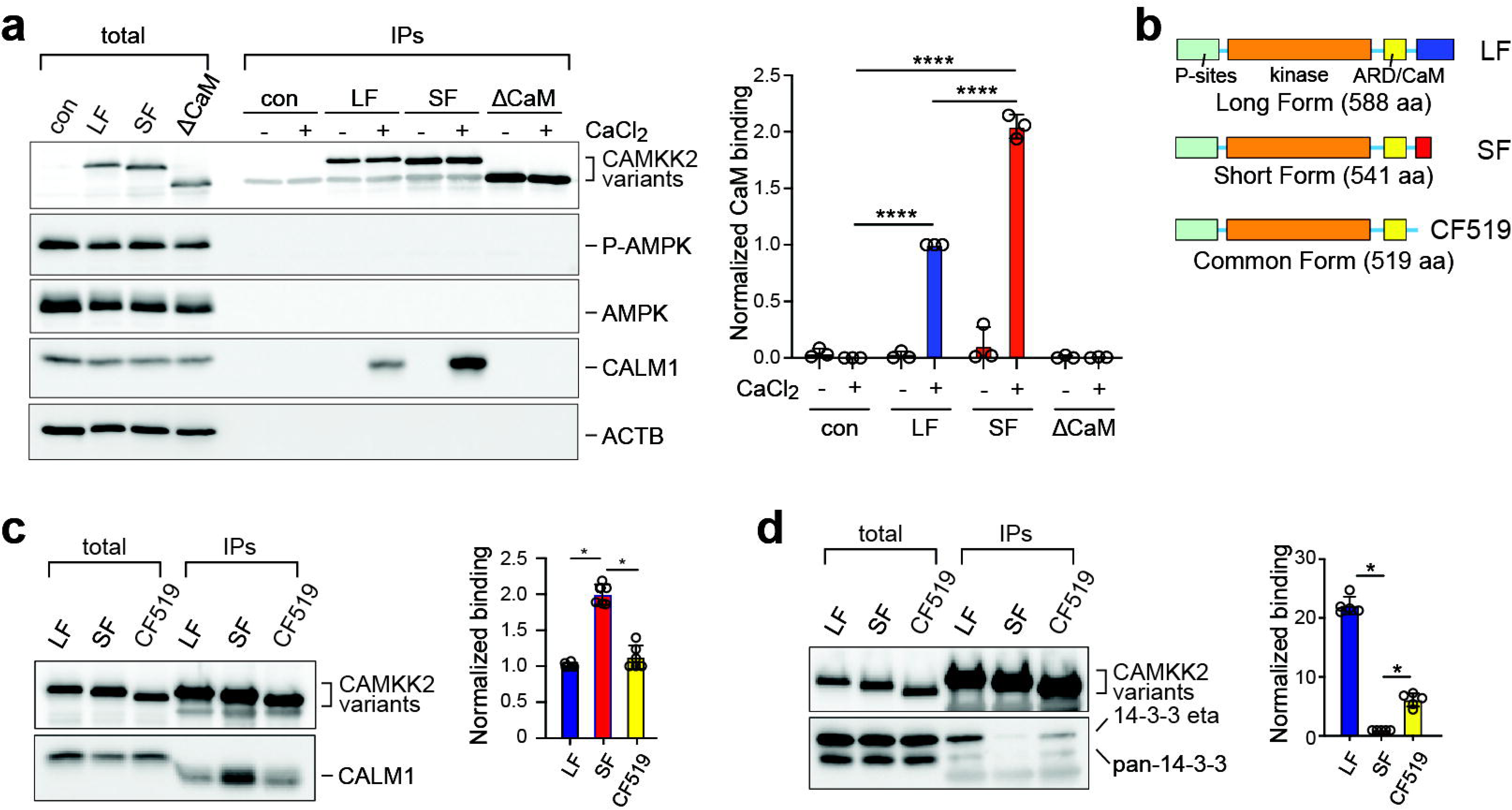
Impact of CAMKK2 isoform-specific C-terminal sequences on protein interactions. **a)** Differential binding of CAMKK2 isoforms to Ca2+/calmodulin. Transiently-transfected Flag-tagged CAMKK2-LF, -SF, and ΔCaM (C-terminally truncated CAMKK2 lacking the auto-inhibitory/Calmodulin binding domain) (471aa) in HEK293 cells were tested for CaM binding using co-IP and western blot analysis. Tubulin was used as a loading control. Quantification of CaM bound (n=3) was determined using densitometry analysis normalized by IP efficiency. Statistical significance was determined by one-way ANOVA followed by Tukey’s post-hoc test from three independent experiments. Significance was denoted by * p-value <0.05, ** p-value <0.01, *** p-value <0.001, **** p-value < 0.0001. **b)** A schematic diagram for the C-terminal protein sequence difference among LF, SF, and the common form 519 (CF519). **c)** CaM co-IP of the transiently-transfected Flag-tagged LF, SF, and CF519 of CAMKK2 in HEK293. **d)** 14-3-3 co-IP of the transiently-transfected MYC-tagged 14-3-3η with the Flag-tagged LF, SF, and CF519 of CAMKK2. Quantification and statistical significance were performed similar to panel A.

To further explore the importance of the C-terminal sequences specific to each isoform, we engineered a truncated CAMKK2 variant (common form 519 amino acids; CF519) containing only the sequence common to both isoforms (Fig. 2b). The truncated variant showed significantly reduced CaM binding, akin to CAMKK2-LF, thereby underscoring the functional significance of the unique C-terminal sequences in CAMKK2-SF for CaM interaction (Fig. 2c). These results also indicate that the C-terminal sequence specific to CAMKK2-LF does not significantly contribute to CaM binding.

In addition to CaM, CAMKK2 also interacts with 14-3-3 scaffolding proteins (ref). Although the binding of 14-3-3 proteins to CAMKK2 has been extensively studied in the context of phosphorylation site mapping^33,34^, the influence of CAMKK2’s C-terminal sequences on this interaction remains underexplored. In experiments similar to those conducted for CaM binding, we utilized both CAMKK2 isoforms and the truncated CF519 mutant to investigate their interactions with 14-3-3 proteins. Contrary to the interactions with CaM, we observed that both 14-3-3 eta (YWHAH) and pan 14-3-3 proteins efficiently co-immunoprecipitated with Flag-tagged CAMKK2-LF whereas the CAMKK2-SF isoform displayed markedly reduced binding. Interestingly, the truncated CAMKK2 variant CF519 maintained its association with 14-3-3 proteins, although to a lesser degree than CAMKK2-LF (Fig. 2d). These findings imply that while CAMKK2-SF predominantly associates with CaM and its related pathways, CAMKK2-LF is preferentially involved with 14-3-3 proteins, impacting different biological pathways. Furthermore, these results highlight that variations in the C-terminal sequences induced by alternative splicing of CAMKK2 may play a crucial role in influencing downstream target pathways.

### Characterization of CAMKK2 isoform exclusive cells

In our characterization of CAMKK2 isoforms through transient expression in HEK293 cells, we observed that the ectopic expression of these isoforms did not influence the phosphorylation of AMPK (AMP-dependent protein kinase) at AMPKα Thr 172, a recognized target of CAMKK2 (Fig. 2a). This suggests that the ectopically overexpressed CAMKK2 might not be fully functional, potentially due to overexpression effects or because the presence of LKB1 in these cells obscures CAMKK2’s impact. Since AMPK can be phosphorylated by both LKB1 and CAMKK2, distinguishing the specific contribution of CAMKK2 is challenging in this context. To address this and specifically analyze AMPK phosphorylation mediated by CAMKK2, we utilized the LKB1-null HeLa cervical cancer cell line and generated a CAMKK2 knockout HeLa cell line using CRISPR-Cas 9 (Fig. 3a). We subsequently engineered the knockout cells to express either CAMKK2-LF or CAMKK2-SF under a doxycycline-inducible promoter, aiming to optimize the expression level close to the endogenous one and isolate the role of CAMKK2 in AMPK regulation. LF cells exhibited a lower AMPK activation compared to SF cells (Fig. 3b). This suggests that the CAMKK2 AS isozymes may have a differential basal activity. Additionally, treatment with increasing concentrations of Doxycycline resulted in a proportional increase in CAMKK2 protein level in both LF and SF cells (Fig. 3c). Using wild-type HeLa cells as a reference, we found that a higher expression of LF is needed to reach to a similar level of AMPK activation than SF cells (Fig. 3c). Furthermore, pharmacological inhibition of CAMKK2 isoforms using the CAMKK2 inhibitor STO-609 showed that SF exhibited a significantly lower IC50 than LF with respect to AMPK activation (Fig. 3d), suggesting that the two isozymes interact differently with the inhibitor, resulting in differences in drug potency. Together, these characterizations suggest that LF and SF are biochemically distinct with respect to substrate activation and inhibitor sensitivity.

**Figure 3.**
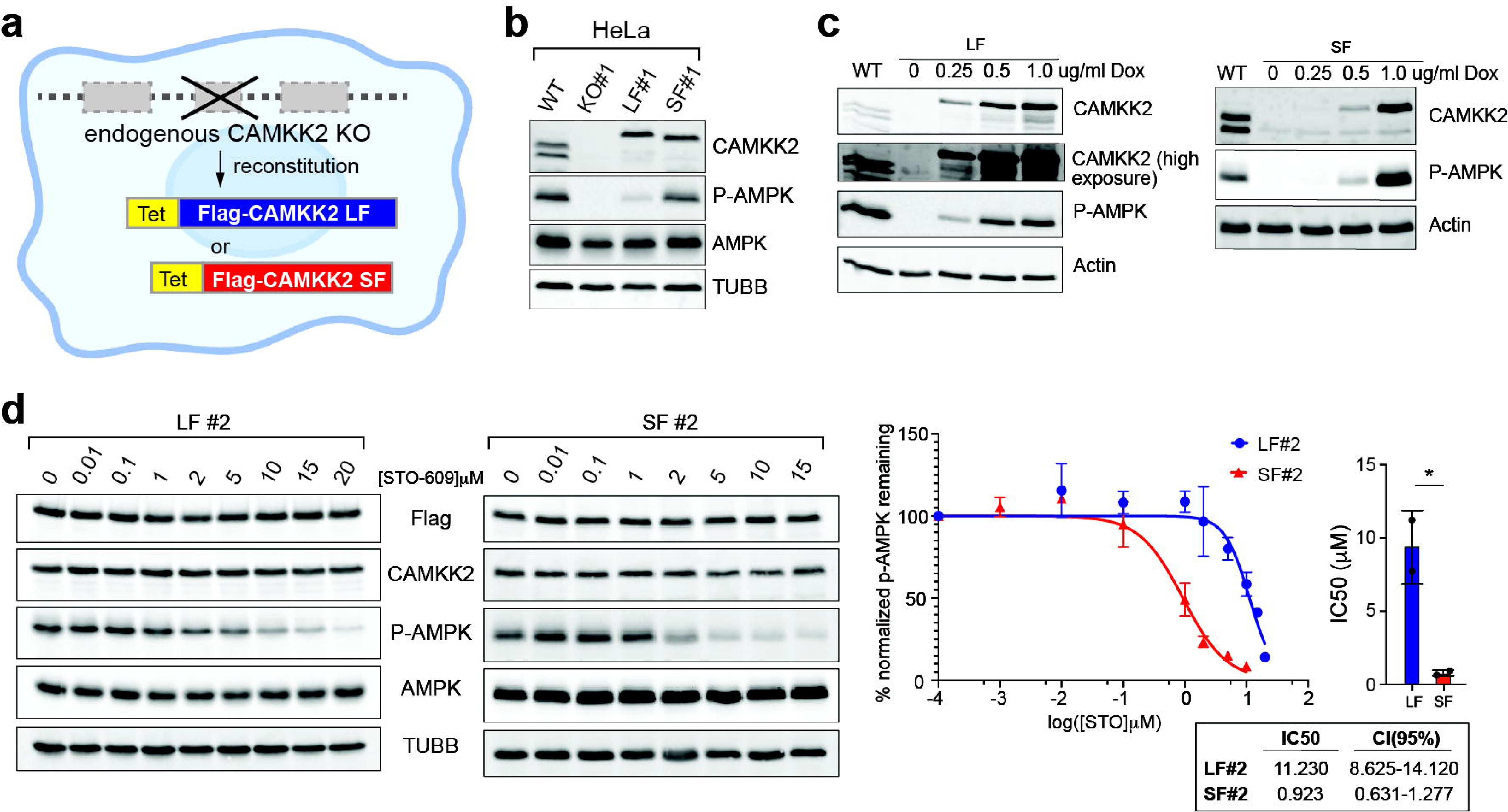
Characterization of CAMKK2 isoform exclusive cells. **a)** Schematic diagram for the engineering of CAMKK2 isoform-specific cell lines using CRISPR/Cas9 gene editing and stable doxycycline-inducible gene integration with drug selection. **b)** Characterization of engineered CAMKK2 isoform-specific HeLa cell lines. The parental wild-type (WT) HeLa cells, *CAMKK2*-knockout clone (KO#1), a dox-inducible CAMKK2-LF (LF#1), and a dox-inducible CAMKK2-SF (SF#1) clones were characterized. P-AMPK (AMPK activation) was used as a readout of the CAMKK2 function. **c)** Representative western blots of CAMKK2 and AMPK phosphorylation in the CAMKK2-LF and -SF cells treated with different doxycycline concentrations. **d**) Representative western blots of dox-induced LF#2 and SF#2 cells treated with various concentrations of the CAMKK2 inhibitor STO-609. P-AMPK was quantified using densitometry analysis and normalized to total AMPK and Flag level and plotted against the log-transformed concentration of STO-609 in a dose-response curve. Non-linear regression was performed to calculate the IC50 value and the 95% Confidence Interval (CI 95%) from the dose-response curve. The IC50 for each sample was plotted on the right. Statistical significance was determined by Student’s t-test with significance denoted by * p-value < 0.05.

### CAMKK2 LF sensitizes cells to glucose-starvation induced cell death

We next investigated the functional relevance of CAMKK2 isoforms in a cancer-relevant environment, specifically, a nutrient-limited setting. Since CAMKK2 is known to regulate energy homeostasis through the AMPK signaling axis^20,24^, we aimed to understand the role of CAMKK2 isoforms in conditions with restricted glucose and serum. Under short-term glucose and serum starvation (<48h), all of the cells exhibited similar levels of cell viability (Fig. 4a). Upon prolonged serum starvation incubation (>68h), KO cells experienced significantly more cell death compared to WT cells, which is consistent with the previously established role of CAMKK2 in energy metabolism (Fig. 4a). Notably, LF cells showed a similar level of cell death to KO cells while SF cells effectively rescued the defects seen in KO cells back to a similar level to that of WT cells (Fig. 4a). These findings confirm that the phenotypic difference seen between LF and SF cells was attributed to the expression of CAMKK2 isoforms. Furthermore, it suggests that SF kinase plays a role in safeguarding cells against cell death induced by nutrient stress.

**Figure 4.**
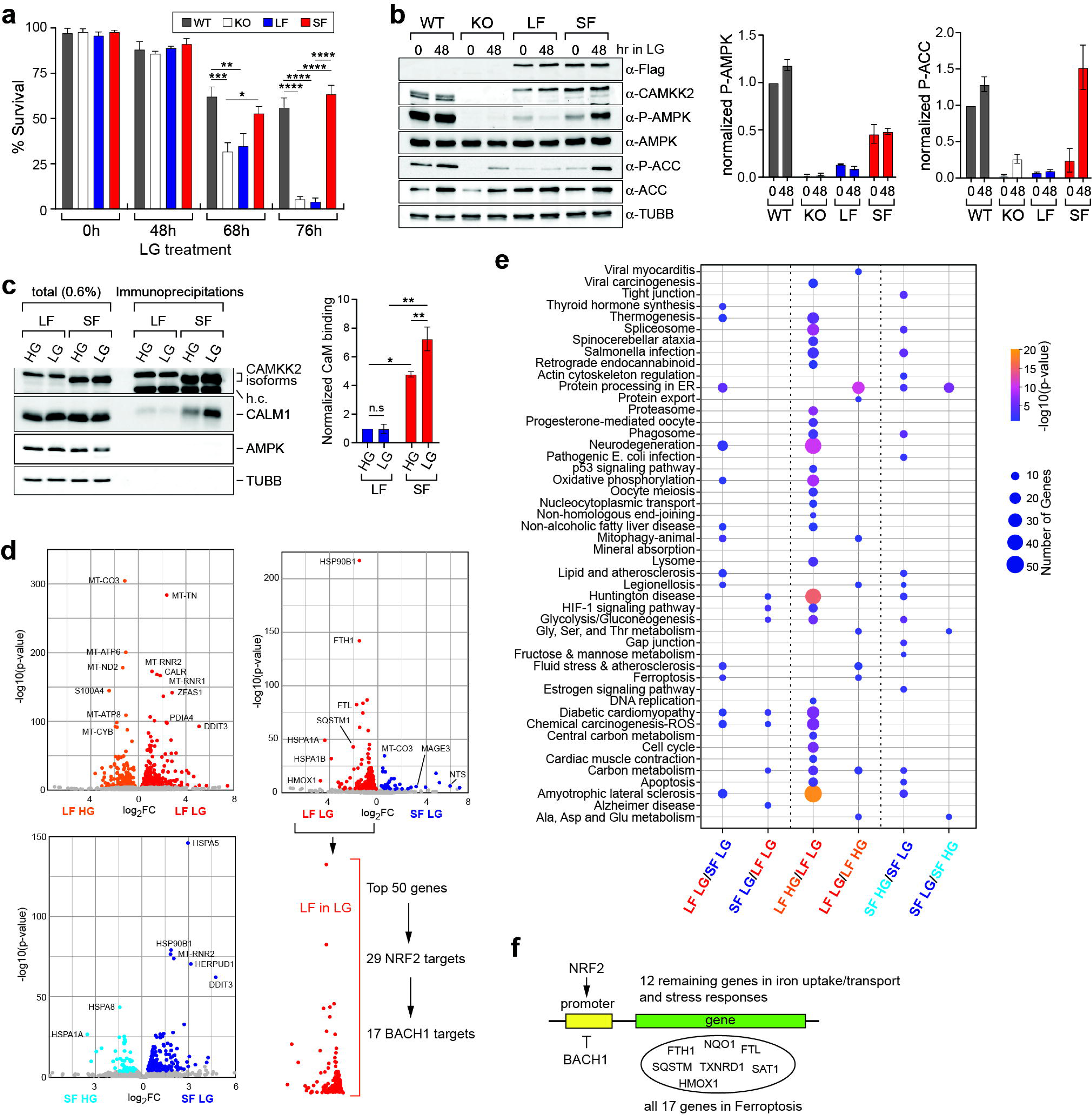
CAMKK2-LF sensitizes cells to glucose-starvation-induced cell death. **a)** Cell viability assay of WT, KO, LF, and SF cells exposed to different durations of glucose starvation. (n=3). % Survival was calculated based on the number of live cells/total number of cells count using Trypan-Blue staining. One-way ANOVA was used to determine the statistical significance at each time point using data from three independent experiments. * p-value < 0.05, ** p-value <0.01, *** p-value <0.001, **** p-value < 0.0001. **b)** Representative western blots of AMPK downstream activation upon glucose starvation in WT, KO, LF, and SF cells. Quantification of the band intensity was performed using densitometry analysis and normalized to total AMPK or total ACC and Tubulin. **c)** CaM co-IP in the control and glucose starved LF and SF cells. **d)** Volcano plots of the differentially expressed genes in the glucose-starved LF vs SF cells, the control vs glucose-starved LF cells, and the control vs glucose-starved SF cells. Differentially expressed genes with statistical significance (p-value<0.01) are color-coded with gray-colored genes denoting no significance. **e)** Bubble plot illustrating the KEGG pathway enrichment of the upregulated genes in the glucose-starved LF vs SF cells, the control vs glucose-starved LF cells, and the control vs glucose-starved SF cells. **f)** BACH1 and NRF2 target genes in the top 50 upregulated genes in LF glucose-starved cells compared to SF.

AMPK activation has been demonstrated to protect cells in maintaining redox control and mitigating oxidative stress upon glucose starvation^35,36^. To elucidate the differential stress responses of the two isozymes, we examined AMPK activity in LF and SF cells during short-term glucose deprivation. Western blot analysis revealed that SF cells either sustained or further enhanced AMPK activation in glucose starved (LG) sample compared to the glucose-rich (HG) sample (Fig. 4b). This pattern of AMPK activation was consistent with WT cells. However, LF cells showed a decrease in AMPK activation upon starvation. Furthermore, the activation pattern of the AMPK downstream substrate acetyl-CoA carboxylase (ACC) activation pattern mirrored AMPK activation in WT, LF, and SF cells, indicating that the AMPK signaling pathway was differentially affected between LF and SF cells upon glucose starvation. Interestingly, while KO cells lack LKB1 and CAMKK2, ACC activation upon glucose starvation was observed, suggesting that KO cells may possess an AMPK-independent pathway for ACC activation. Together, glucose starvation treatment unveiled the metabolic vulnerabilities of LF cells as well as the inability of LF kinase to activate the stress-protective AMPK pathway in response to nutrient stress. AMPK activation by CAMKK2 requires Calmodulin binding^20^. To delineate how the differential AMPK activation was observed in glucose-starvation between CAMKK2 isoforms, we hypothesized that it is due to the extent of calmodulin binding. Flag-immunoprecipitation of doxycycline-induced CAMKK2 isoforms upon starvation showed that calmodulin binding remained defective for LF kinase, but SF kinase showed a further increase in binding to calmodulin upon starvation, consistent with the increased or sustained high AMPK activation seen in SF cells (Fig. 4c).

Besides AMPK signaling, we further assessed the transcriptome-wide impact of LF and SF with RNA-sequencing to quantify global gene expression changes in LF and SF cells experiencing glucose starvation. Interestingly, when comparing between the glucose-rich (control) and the glucose-starved conditions within both LF and SF cells, we observed a total of 755 and 274 differential expressed genes, respectively (Fig. 4d). The high number of differential expressed genes found in LF control and glucose-starved sample suggests a more extensive reprogramming of the transcriptome in LF cells compared to SF cells. The KEGG pathway analyses of these differentially expressed genes showed the enrichment of several metabolic pathways including Glycolysis/Gluconeogenesis, Carbon Metabolism, Glycine/Serine/Threonine metabolism, and Alanine/Aspartate/Glutamate metabolism common to both LF and SF cells (Fig. 4e). These common pathways suggest that glucose level can dynamically induce changes in the transcriptome landscape independent of CAMKK2 isoform expression. To discern the transcriptomic difference between LF and SF cells under glucose starvation, KEGG analysis of the differential expressed genes directly comparing glucose-starved LF and glucose-starved SF showed interesting pathway enrichment differences (Fig. 4e). For instance, pathways including HIF-1 signaling and biosynthesis of amino acids were found to be enriched in upregulated genes in glucose-starved SF whereas genes involved in oxidative phosphorylation, ferroptosis, mitophagy and lipid and atherosclerosis were upregulated in glucose-starved LF cells. To further understand how the transcriptomic changes by glucose starvation might lead to the cell death observed, we selected the 50 most upregulated genes in LF cells comparing to SF cells upon glucose starvation and conducted a transcription factor network analysis using TFLINK^37^, a known human transcription factor database, to determine mechanistically how the transcriptome was regulated in LF compared to SF glucose-starved cells. Among the 1051 transcription factors shown to interact with or have binding sites on the top 50 genes in LF glucose-starved cells, we found NRF2 and BACH1 to have prominent control of these genes (Fig. 4f). These NRF2 and BACH1 target genes with an upregulation in LF glucose starved cells were enriched in several cellular stress and cell death pathways were highlighted, such as ferroptosis, necroptosis, and fluid shear stress and atherosclerosis in LF cells (Fig. 4g). This further reveals the stress levels LF cells exhibited and matches with the higher cell death observed in LF cells upon starvation. Together, our findings suggest NRF2 and BACH1 transcriptional control within the LF and SF transcriptome during glucose starvation.

### Glucose starvation induces oxidative stress in LF cells, activating NRF2/HO1 ferroptosis cell death

Glucose starvation has been shown to induce ferroptosis, a form of non-apoptotic regulated cell death dependent on iron accumulation and lipid peroxidation^38,39^. As enriched as the top KEGG pathway, we sought to analyze the expression of more than 60 ferroptosis-related genes taken from the Gene Set Enrichment Analysis (GSEA) database. Interestingly, when comparing the glucose-starved LF and SF cells, the heatmap showed opposing, differential expression of approximately half of the ferroptosis-related genes (visualized on the left/top heatmap) while the other half of the genes exhibited similar expression between the cells (visualized on the right/bottom heatmap) (Fig. 5a). The selected upregulated ferroptosis-related genes in LF cells are known as pro-ferroptosis genes while the anti-ferroptosis genes were upregulated in SF cells (Fig. 5a). This transcriptomic pattern suggests that glucose starvation induces a unique ferroptotic signature in LF cells with a focus in *HMOX1* induction. We examined Heme oxygenase 1 (HO-1) protein level in the control and glucose-starved WT, KO, LF, and SF cells and found a consistent reduction of HO-1 expression in glucose-starved WT and SF cells and an increased expression in KO and LF cells similar to the RNA-seq results (Fig. 5b). These results together suggest that reconstituted LF cells remain sensitive to ferroptosis upon starvation whereas reconstituted SF cells suppress ferroptosis, similarly to the WT cells. To understand how the transcription of *HMOX1* was selectively promoted in LF cells, we assessed the level and nuclear translocation of transcription factors NRF2 and BACH1, which are known to regulate *HMOX1* expression. Interestingly, LF and SF cells showed different transcriptional regulation of *HMOX*1. Specifically, BACH1 was selectively upregulated in SF with an increase in nuclear translocation upon starvation. On the other hand, NRF2 was selectively upregulated and translocated to the nucleus in LF cells (Fig. 5c). BACH1 and NRF2 are known to serve opposing roles in *HMOX1* transcription^40–43^. To understand the relationship between BACH1 and NRF2 on *HMOX1* transcription, we performed a siRNA knockdown of BACH1, NRF2, and a double knockdown and assessed how the knockdown would affect *HMOX1* expression. We found that the knockdown of NRF2 didn’t change *HMOX1* expression compared to the control knockdown (Fig. 5d). However, upon BACH1 knockdown, *HMOX1* expression was increased significantly compared to the control knockdown. When BACH1 and NRF2 were both depleted, *HMOX1* expression was suppressed compared to BACH1 knockdown alone. This suggests that BACH1 plays a major suppressive role in *HMOX1* transcription. Upon BACH1 dissociation from the *HMOX1* promoter, NRF2 promotes *HMOX1* transcription.

**Figure 5.**
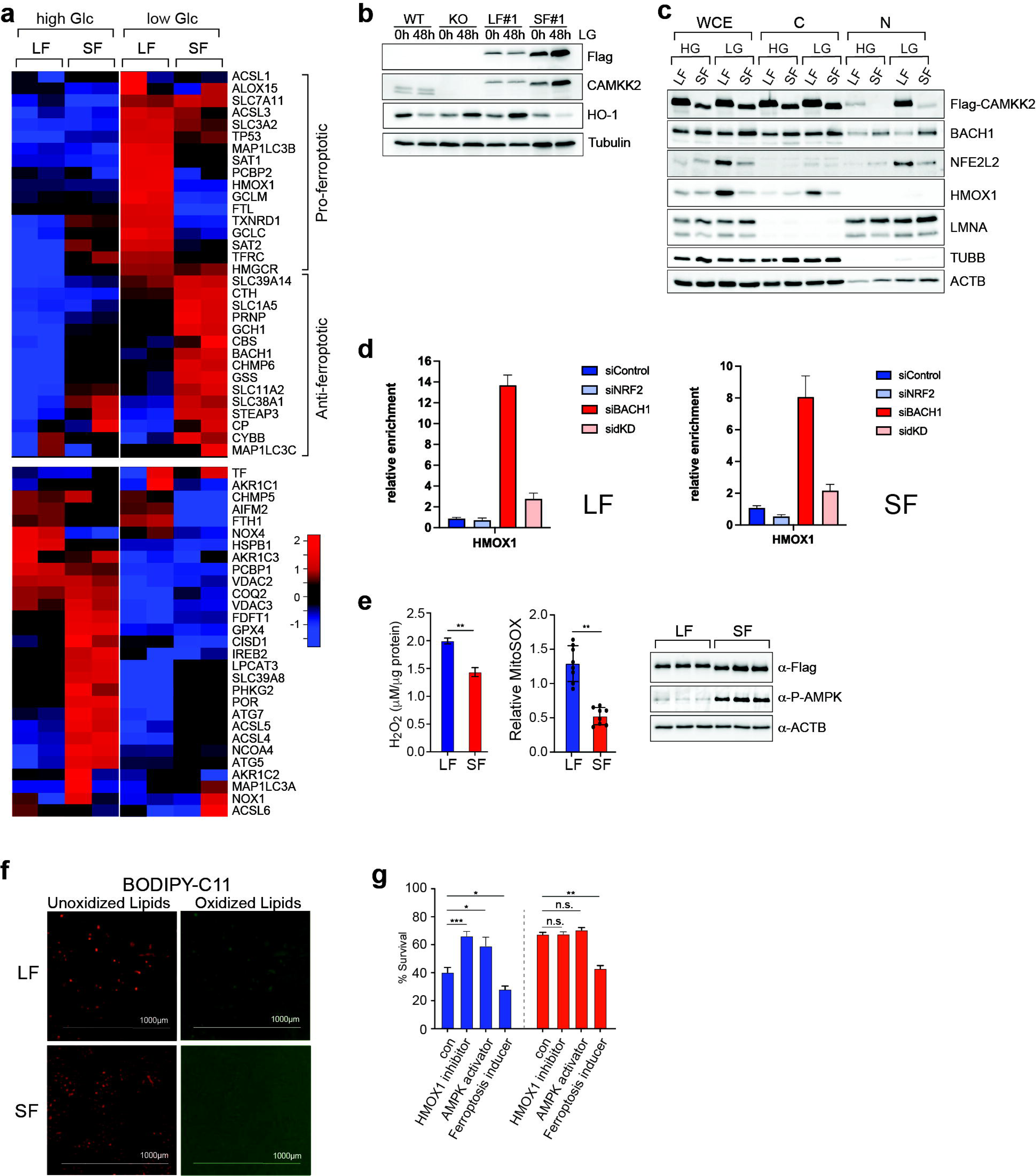
Glucose starvation induces oxidative stress in LF cells, activating NRF2/HO1 ferroptosis cell death. **a)** Hierarchical clustering and heatmap of expression data of ferroptosis-related genes in the control and glucose-starved LF and SF RNA-sequencing datasets (n=2). The list of ferroptosis genes was taken from the Gene Set Enrichment Analysis (GSEA) database. **b)** Representative western blots of heme-oxygenase (HO-1) protein level in control and glucose-starved WT, KO, LF, and SF cells. **c)** Representative western blots of BACH1 and NRF2 in whole cell lysates, nuclear fraction, and cytoplasmic fraction of control and glucose-starved LF and SF cells. **d)** qPCR of *HMOX1* expression in the siControl, siBACH1, siNRF2, and double siBACH1/siNRF2 LF and SF cells. **e)** Hydrogen peroxide assay and mitoSOX measurement in glucose-starved LF and SF cells. (n>3). **f)** Representative images of BODIPY-C11 staining in glucose-starved LF and SF cells (n>3). **g)** Cell viability assay of glucose-starved LF and SF cells treated with vehicle control, ZnPP, or A769662. (n=3). % Survival was calculated based on the number of live cells/total number of cells count using Trypan-Blue staining. One-way ANOVA was used to determine the statistical significance at each time point using data from three independent experiments. * p-value ** p-value <0.01.

Reactive oxygen species (ROS) produced during glucose starvation-induced cellular stress promote ferroptosis by increasing lipid peroxidation^35,44,45^. To test whether LF and SF cells experienced differential cellular stress during glucose starvation, cellular ROS level was measured. LF cells showed higher levels of ROS compared to SF cells upon starvation (Fig. 5e). Additionally, LF cells also showed higher lipid peroxidation, as observed with a higher fluorescence intensity by the lipid peroxidation fluorescence dye BODIPY C11 (Fig. 5f). This supports that LF and SF cells confer different sensitivity to oxidative stress and ferroptosis upon glucose starvation. To determine if HO-1 upregulation or AMPK inactivation resulted in the LF cell death in glucose starvation, we treated HMOX1 inhibitor (ZnPP) and AMPK activator (A769662) to LF and SF cells in glucose starvation media and determined the cell death. We found that both ZnPP and A769662 treatment to LF cells rescued LF the cell death similar to SF level without further benefiting the viability of SF cells (Fig. 5g). This suggests that both AMPK inactivation and HMOX1 upregulation mechanistically sensitize LF cells to glucose starvation. Together, these findings demonstrated the differential novel functions of the alternatively spliced isoforms of CAMKK2 in energy metabolism. The defect in AMPK activation and the upregulation of HMOX1 by the CAMKK2 LF isoform induces ferroptosis in a nutrient-limited environment in cancer cells (Fig. 6).

**Figure 6.**
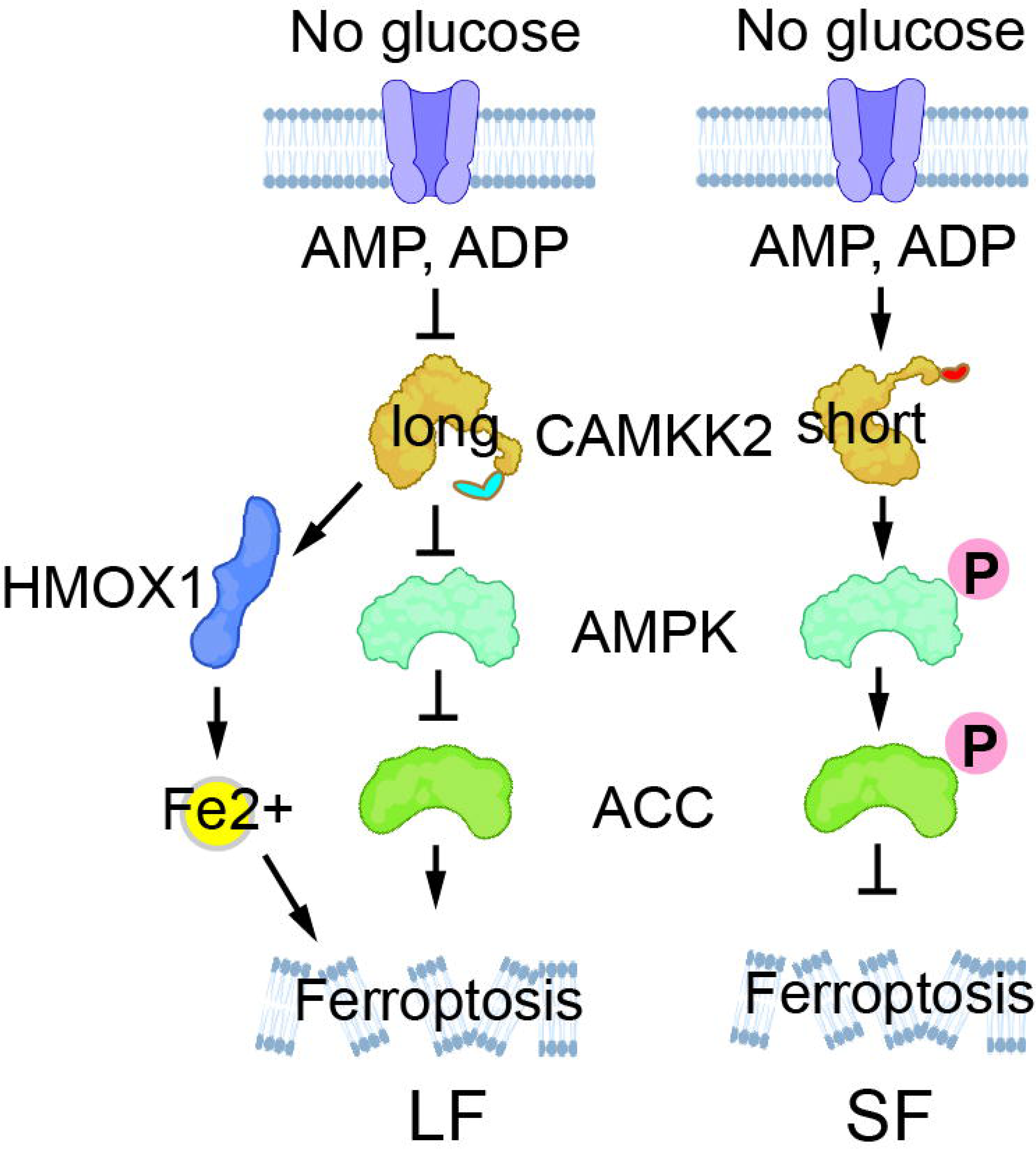
Schematic summary model of the differential role of CAMKK2 isoforms during glucose deprivation. During glucose-starved conditions, CAMKK2 SF induces downstream AMPK signaling, thereby inhibiting ferroptosis and metabolic stress-induced cell death. Conversely, CAMKK2 LF fails to activate AMPK signaling and in turn, upregulates heme oxygenase-1 (*HMOX1/*HO-1) levels, promoting ferroptosis and cell death.

## Discussion

The emergence of mTOR signaling regulation in alternative processing of RNAs expands its multifaceted roles in gene expression regulation, in addition to its well-documented functions in protein translation, autophagy, and metabolism^46^. Specifically, several mechanistic studies supported the link of mTOR signaling to 3’-UTR shortening through alternative polyadenylation^13,47^. However, the functional impact of mTOR’s regulation on alternative splicing remains understudied. A previous study discovered that mTORC1-mediated phosphorylation of RNA-binding protein SRPK2 enables its association with spliceosomal protein U1-70K, promoting splicing and suppressing intron retention of pre-mRNAs for genes involved in lipogenesis^48^. Global profiling of alternative splicing in cells with mTOR inhibition also revealed transcriptome-wide exon skipping^49^. These studies collectively suggest the role of mTOR signaling in alternative splicing, although the functional consequences of these AS changes are unknown.

Our study unveiled the cancer-relevant *CAMKK2* AS isoform switch mediated by mTOR signaling through the upregulation of splicing factor SRSF3. We found that the C-terminally distinct CAMKK2 AS isozymes are biochemically distinct with its protein binding preference. Additionally, the differences we observed in the STO-609 IC50 value for CAMKK2 isozymes further suggest that the unique disordered C-terminus of these isozymes plays a significant role in regulating the kinase’s activity. While both CaM and 14-3-3 have been shown to interact with CAMKK2^33,34^, the isoform-specific preference in protein binding suggests that the C-terminal sequences of CAMKK2 produced by alternative processing of mRNA possess roles in influencing protein-protein interactions and regulate downstream CaM and 14-3-3 pathways. Previous CAMKK2 structural insights are solely from the soluble portion of CAMKK2 lacking the unique C-terminal sequences, it is unknown how the tail might interact with known CaM and 14-3-3 binding regions. Future studies will focus on how the C-terminus of CAMKK2 isoforms contributes to protein-protein interaction by identifying individual critical residues in the unique region of the isoforms through serial tail deletion and assess CaM and 14-3-3 binding. There are also several unique phosphorylation and other post-translational modification sites found on the C-terminus that can potentially affect the CAMKK2 kinase function^20,29^. Further biochemical and structural characterization of the C-terminal tail of the isozymes with Surface Plasmon Resonance (SPR) assay to determine the detailed biophysical properties of CAMKK2 to CaM and 14-3-3.

Glucose starvation is commonly found in cancer cells, which rely on AMPK pathway activation to inhibit anabolic pathways and promote survival during stress conditions like hypoxia and nutrient deprivatione^35,50^. LKB1, the primary upstream kinase responsible for AMPK activation in cancer cellsl^21,50^, is frequently found to be non-functional by loss-of-function mutations in 20-30% of cervical and lung cancers^51,52^. Patients with LKB1-mutant cancer patients with LKB1 mutations exhibit chemotherapy resistance, higher metastasis risk, and worse overall survival compared to patients with wild-type LKB1. In HeLa cells (a LKB1 deficient cervical cancer cell line), we uncovered that the SF of CAMKK2 serves as the sole kinase for AMPK activation. HeLa cells expressing LF of CAMKK2 or lacking CAMKK2 showed disrupted AMPK signaling and increased ferroptotic stress, leading to cell death under glucose starvation. This explains why cancer cell lines and patient tumor samples exclusively expresses CAMKK2 SF as an adaption to enhance metabolic fitness. Further investigations into mechanisms driving CAMKK2 exon inclusion could identify novel therapeutic targets for cancer treatments.

Our study has established a novel axis involving CAMKK2-AS/NRF2-BACH1/HMOX1 in the regulation of ferroptosis. Ferroptosis is an emerging target for therapeutic intervention in cancer and inflammatory diseases^53,53,54^. Various genes involved in oxidative stress, lipid peroxidation, and their transcription factors have been linked to ferroptosis. However, many of these genes have been reported to be both pro- and anti-ferroptotic^55–57^, thus their roles in ferroptosis are highly context-dependent. In our glucose starvation RNA-sequencing data, the 4-fold upregulation of HMOX1 in LF cells upon starvation points to a specific mechanism triggering ferroptosis through NRF2/BACH1/HMOX1 pathway. Further investigation into how ferroptosis is triggered in LF cells during starvation particularly whether the involvement of CaM and AMPK signaling is necessary to determine the exact mechanism.

In this study, we provided evidence demonstrating the biochemical and functional significance of CAMKK2 alternative splicing isoforms. Our genome-wide analysis revealed that approximately 48% of human kinase genes undergo alternative splicing near the terminal coding exon, generating kinase isoforms with conserved functional domains but variable C-termini. With the vast majority of these annotated isoforms still unexplored, our study highlights the urgent need to delineate the functions and pathological relevance of these widespread C-terminal proteoforms.

## Methods

### Cell lines and cell culture

WT MEF and *Tsc1-/-* MEFs were obtained from Dr. Kwiakowski laboratory at Harvard University and were previously published [PMID: 11875047]. WT MEFs, *Tsc1-/-* MEFs, HeLa, HEK293, HCT116, A549, U2OS, MDA-MB231 cells were cultured in High Glucose (4.5g/L) Dulbecco’s Modified Eagle Media (DMEM) (Gibco), supplemented with 10% (v/v) fetal bovine serum (FBS) (Gibco), 100ug/ml Penicillin-Streptomycin at 37°C with 5% CO_2_. Growth factor and amino acid starvation experiments in HeLa cells were done using Earle’s Balanced Salt Solution (EBSS) (Gibco). Glucose starvation experiments were conducted using Low Glucose (1g/L) Dulbecco’s Modified Eagle Media (DMEM) (Gibco), in supplements with 1% dialyzed FBS (Sigma).

### Cloning of CRISPR-Cas9-sgRNA plasmids and generation of CAMKK2 knockout cells

The gRNA sequences were identified and designed by crispor.tefor.net. The guides were cloned into PX458 plasmid (Addgene #48138) using the following primer pairs. Guides targeting intron 8 region were forward primer 5’- CACCGTAGCTGAGTGAGGTGCCACT -3’ and reverse primer 5’- AAACAGTGGCACCTCACTCAGCTAC-3’. Guides targeting intron 9 region were forward primer 5’- CACCGCAGAATGCTAACTACCCACT-3’ and reverse primer 5’- AAACAGTGGGTAGTTAGCATTCTGC-3’. The plasmids were co-transfected into WT HeLa cells via electroporation (Neon Transfection System, Thermo Fisher Scientific) for 48hrs prior to sub-cloning in 96-well plates. Individual clones were screened with genomic DNA-templated PCR using forward primer 5’- TGTTCGAACTGGTCAACCAAGG-3’ and reverse primer 5’- CTGTGCTCCAGGAAGAAAATAAG-3’ and confirmed by Western blot analysis.

### Cloning and generation of dox-inducible or Flag-tagged constitutive CAMKK2 isoform-specific cell lines

Human CAMKK2 SF was first amplified with cDNA template from HeLa cells using the primers For: GACAAGCTTTCATCATGTGTCTCTAGCCAGCCCAGCAGC and Rev:TAAGGATCCCTACTCGGGCTCCATGGCCTCCTCCGGC. The PCR with the correct expected size for SF was purified and digested with HindIII and BamHI and ligated into p3X-Flag-CMV10 plasmid. Using the successfully cloned CAMKK2 SF containing plasmids, the cloning of CAMKK2 LF and CAMKK2 (ΔAID/CaM) was subsequently performed using inverse PCR method with:

LF forward primer: AGTCCCTGTCTGAGCTCAAGGAAGCAAGGCAGCGAAGACAACCTCCAG
and LF reverse primer: CACATTCCCTGGTTGGTTTTTTGGTGAGCAAGTTTCCAGGCGCTGACAG
or ΔAID/CaM forward primer: TAGAACTCAGTCAAACACATTCCCAGC ΔAID/CaM
reverse primer: GACCTCCTCTTCAGTCACTTCGACCAG.
The cloning of CAMKK2 CF519 was performed with forward primer: TAGGGATCCCGGGTGGCATCCCTGTGAC
and reverse primer: TTTGGTGAGCAAGTTTCCAGGCGCTGACAGTGAG.

The cloning of the Flag-tagged LF and SF into pCW57.1 plasmid was performed using the Flag-tagged LF and SF-inserted-p3X-Flag-CMV10 plasmids as templates to amplify the cDNA region with NheI and SalI and ligated into pCW57.1 plasmid.

Plasmids were transfected into CAMKK2 Knockout HeLa cells using jetPRIME transfection reagents (Polyplus) for 48 hours and selected using 1ug/ml puromycin to generate stable cell lines. The stable cell pools were sub-cloned in 96-wells and screened using Western blot analysis in the presence and absence of doxycycline.

### Chemicals

STO-609 (No. 52029-86-4), Torin 1 (No. 10997), doxycycline (No. 14422), ZnPPIX (No. 14483), A769662 (No. 11900), A23187 (No. 11016), were purchased from Cayman Chemical (No. 52029-86-4) and reconstituted in DMSO.

### RNAi knockdown

Cells were transfected with siRNAs (Integrated DNA Technologies) using RNAiMax Transfection reagents (Thermo Fisher Scientific) recommended by the manufacturer’s protocol. siRNAs were designed and synthesized by IDT targeting the following mRNA regions:

mSrsf1: UGGAAUUUGAAGGUAAGGAUAC,
mSrsf2: GGUCUAUAUGAUAAUCUGAACCCAA,
mSrsf3: CGUGAUAUCAAGAAUGUUACUUUA,
mSrsf4: GCAGUGGAUAUGGUUAUCGAAGAAG,
mSrsf5: AUCCGAGAUAUUGACUUGAAAAGAG,
mSrsf6: CAUGAAGCGUGCUUUGGAUAAACUG,
mSrsf7: GAGUGAGGGUUGAACUAUCAACAGG,
mSrsf9: CUAUGGAAGAAACGGUUACGAUUAU,
mSrsf10: GAAGACGCUUUACAUAAUUUGGACA,
mSrsf11: GCUAUUGAAACUUAUGAGUACUGUU,
hBACH1: GAGGAAUCCUGCUUUCAGUUUCUGA,
hNFE2L2: AUUGAUGUUUCUGAUCUAUCACUTT

### Co-immunoprecipitation (co-IP)

For CaM co-IP, Flag-CAMKK2 transfected HEK293 cell pellets were resuspended in lysis buffer (50mM Tris-HCl pH7.5, 150mM NaCl, 0.2% Tx100, 1mM EDTA, and 1mM EGTA supplemented with protease and phosphatase inhibitor cocktails. After sonication and centrifugation, CaCl_2_ (final concentration 3mM) was added to the supernatants containing 300-500ug of total proteins before immunoprecipitation using anti-FLAG magnetic beads (M8823, Millipore Sigma) for 2 hours at 4°C. Beads were washed with lysis buffer 5 times. The immunoprecipitated proteins were eluted using 2x Laemmli Sample Buffer and run on SDS-PAGE gels for western blot analysis. For 14-3-3 co-IP, Flag-CAMKK2 and MYC-14-3-3η (a gift from Dr. Andrey Shaw, Washington University, St. Louis) plasmids were co-transfected into HEK293. Cell pellets were resuspended in a milder lysis buffer (20mM Tris-HCl pH7.5, 70mM NaCl, 0.01% Tx100 supplemented with protease and phosphatase inhibitor cocktails) and the immunoprecipitation took place for 3 hours.

### Cell fractionation and nuclear enrichment

Subcellular fractions were extracted using the method from Suzuki et al. 2010 (PMID: 21067583). Briefly, cell pellets were resuspended in lysis buffer (25mM Tris-HCl pH7.5, 100mM NaCl, 0.1% NP-40 supplemented with protease and phosphatase inhibitor cocktails). A portion of the resuspended cells was saved as the whole cell lysate. The rest of the resuspended cells were subjected to quick (10 seconds), low-speed centrifugation. Supernatants containing the cytoplasmic fraction were collected. Pellets containing the nuclear fraction were washed 5 times with lysis buffer and resuspended in 2x Laemmi Sample Buffer. Whole cell lysates, cytoplasmic fractions, and nuclear fractions were run on SDS-PAGE gels for western blot analysis.

### Western blot analysis

Harvested cell pellets were lysed with ice-cold 25mM Tris-HCl pH 7.5, 100mM NaCl, 0.05% Tx100 supplemented with protease and phosphatase inhibitor cocktails (Sigma) and sonicated with an ultrasonic processor (AE Bios) with manufacturer’s recommendation. Supernatants were collected after centrifugation at 12,000 rpm for 10 min at 4°C. Protein concentration was determined using the Pierce BCA Protein assay (Thermo Fisher Scientific). Lysates with equal ug of proteins were loaded and run on SDS-PAGE and transferred to nitrocellulose membrane. Blots were blocked with 5% BSA in TBST. Antibodies used for western blot analysis include anti-Flag (No. F3165, Sigma Aldrich), anti-Actin (612657, BD Biosciences), anti-CAMKK2 (sc-100364, Santa Cruz Biotechnology), anti-phospho-AMPK (No. 2535, Cell Signaling), anti-AMPK (No. 5831, Cell Signaling), anti-phospho-ACC (No. 11818, Cell Signaling), anti-ACC (No. 3676, Cell Signaling), anti-Tubulin-Rhodamine (12004166, Bio-Rad), anti-Calmodulin (No. 35944, Cell Signaling), anti-HO1 (No. 43966, Cell Signaling), anti-BACH (14018-1-AP, Proteintech), anti-NRF2 (No. 12721, Cell Signaling), anti-14-3-3 (pan) (No. 8312, Cell Signaling). Anti-Secondary antibody against mouse: goat-anti-mouse IgG-HRP (No. sc-2005, Santa Cruz Biotechnology), Secondary antibody against rabbit: goat-anti-rabbit IgG-HRP (No. sc-2004, Santa Cruz Biotechnology), StarBright Blue 700 Goat Anti-Mouse IgG (12004159, Bio-Rad). Western blot images were captured with the ChemiDoc MP Imaging System (Bio-Rad).

### Real-time PCR (RT-PCR) and Real-time quantitative PCR (RT-PCR) analysis

Total RNAs were extracted using the Trizol method recommended by the manufacturer’s protocol. cDNAs were made by reverse transcription using Oligo-d(T) and superscript III (Thermo Fisher Scientific). SYBR Green fluorescence was used for quantitative real-time PCR reactions. The relative expression between groups was measured using the ΔΔCt method.

The list of primers used for RT-PCR and qRT-PCR is:

mouse *Camkk2* RT-PCR forward: 5’-GGGAACCCGTTCGAAGGTAG-3’;
mouse *Camkk2* RT-PCR reverse: 5’-GCAGGGACCACCTTTCACAA-3’;
human *CAMKK2* RT-PCR forward: 5’-ACGTAAACGCTCCTTTGGGA-3’;
human *CAMKK2* RT-PCR reverse: 5’-AACTTTCCACGCAGGGACTA-3’;
human *CAMKK2* qPCR forward: 5’-GGATGACCCCAATGAGGACC-3’;
human *CAMKK2* qPCR reverse: 5’-GGAAGTAGAAACGGGCCTGG-3’;
human *HMOX1* qPCR forward: 5’-CAACAAAGTGCAAGATTCTGCCCCC-3’;
human *HMOX1* qPCR reverse: 5’-TGTCGCCACCAGAAAGCTGA-3’;
human *LCN2* qPCR forward: 5’-GGACAACCAATTCCAGGGGA-3’;
human *LCN2* qPCR reverse: 5’-TTCAGCTCATAGATGGTGGCA-3’;
human *HSPA1A* qPCR forward: 5’-GGGCCTTTCCAAGATTGCTG-3’;
human *HSPA1A* qPCR reverse: 5’-TGCAAACACAGGAAATTGAGAACT-3’;
human *HSPA1B* qPCR forward: 5’-CAGGCCCTACCATTGAGGAG-3’;
human *HSPA1B* qPCR reverse: 5’-CAGCAAAGTCCTTGAGTCCCA-3’;
human *GPX4* qPCR forward: 5’-GGACAAGTACCGGGGCTTCGTG-3’;
human *GPX4* qPCR reverse: 5’-ACTCAGCGTATCGGGCGTG-3’;
human *SLC7A11* qPCR forward: 5’-TGCAGTGGCAGTGACCTTTTCTGA-3’;
human *SLC7A11* qPCR reverse: 5’-ACCACCGTTCATGGAGCCAA-3’;
human *UBE2B* qPCR forward: 5’-CGGTTACAAGAGGACCCACC-3’;
human *UBE2B* qPCR reverse: 5’-GGTGTCCCTTCTGGTCCAAAT-3’;
human *NFE2L2* qPCR forward: 5’-AACCAGTGGATCTGCCAACT-3’;
human *NFE2L2* qPCR reverse: 5’-GGGAATGTCTGCGCCAAAAG-3’;
human *FTL* qPCR forward: 5’-ACCTCTCTCTGGGCTTCTATTTCG-3’;
human *FTL* qPCR reverse: 5’-GCATCTTCAGGAGACGCTCG-3’;
human *FTH* qPCR forward: 5’-CGGGCTGAATGCAATGGAGT-3’;
human *FTH* qPCR reverse: 5’-ATGAAGTCACACAAATGGGGGTCATT-3’;
human *NQO1* qPCR forward: 5’-GTACTGGCTCACTCAGAGAGGAC-3’;
human *NQO1* qPCR reverse: 5’-ATGGCATAGAGGTCCGACTCCACCA-3’;
human *BACH1* qPCR forward: 5’-AGACGACTCTGAGACGGACA-3’;
and human *BACH1* qPCR reverse: 5’-AGACAGTGAAATTATCCGTTGTGC-3’.

### Cell Viability Assay

WT, KO, LF, and SF HeLa cell lines were seeded in 6-well plates for 48 hours in High Glucose media to reach to ∼60% confluency. These cell lines were incubated in Low Glucose media for a total of 68 hours. Doxycycline was added in LF and SF cells to induce CAMKK2 isoform expression over the entire course of the assay. Cells absence of doxycycline were used as controls. Over the time course, samples were collected and mixed with equal volumes of 0.4% Trypan Blue staining solution and counted using the Countess II (Thermo Fisher Scientific). % Survival was calculated using the number of live cells over total cells.

### Hydrogen peroxide assay

Hydrogen peroxide (H_2_O_2_) was quantified using the Amplex Red Hydrogen Peroxide/Peroxidase Assay Kit (A22188, Invitrogen) using the manufacturer’s protocol. Cells were lysed in 250mM Sodium Phosphate pH 7.4 with 0.1% Tx100 and sonicated. 50ul of cell extracts were incubated with 50ul of Amplex Red reagent and Horseradish Peroxidase solution for 30min in RT. Fluorescence was measured using a fluorescence microplate reader with an excitation wavelength of 540nm and emission wavelength of 590nm and normalized to protein concentration. A standard curve was used to determine the absolute amount of H_2_O_2_ present in the samples.

### Mitochondrial ROS assay

Mitochondrial ROS levels were measured using the MitoSOX Red superoxide dye (M36008, Invitrogen) according to the manufacturer’s protocol. Live cells were stained with 5uM of MitoSOX Red for 30min, washed, and the fluorescence was measured using a fluorescence microplate reader. The relative fluorescence was normalized to the cell number determined by Hoechst 33342 staining (H1399, Invitrogen).

### Lipid peroxidation Assay

Control and glucose-starved LF and SF HeLa cells were washed with Hanks’ Balanced Salt Solution (HBSS) before staining with 1uM of BODIPY 581/591 C11 (D3861 Invitrogen) fluorescent dye for 30 min at 37C. Stained cells were washed twice with HBSS before imaging with BioTek Cytation C10 Confocal Imaging Reader (Agilent). Fluorescence intensity was quantified and normalized to cell number.

### RNA-seq and Data Analyses

Total RNA was extracted from WT MEFs, Tsc1-/-MEFs, WT HeLa, KO, LF, and SF cells using the Monarch Total RNA Miniprep Kit (New England BioLabs) and submitted to the University of Minnesota Genomics Center (UMGC) for RNA-sequencing analysis. RNA samples was quantified using the RiboGreen Assay and the quality of the RNA was determined by the BioAnalyzer (Agilent) and confirmed to have a minimum of a passing RIN score of 8. Samples were library-prepped and sequenced on the NextSeq 2K for 2×150bp with ≥ 60M reads generated per human samples and ≥ 80M reads per mouse samples. The quality score for all of the libraries was ≥Q30. Paired-end reads were aligned to the human hg38 or mouse mm10 reference genome using TopHat^58^ with a maximum of two mismatches allowed. Kallisto^59^ was applied to quantify gene expression with UCSC annotation^60^. TPM (transcript per million) > 1 was used as the minimum cut-off to be considered as expressed genes. Differential gene expression analysis (DGE) was performed using DESeq2^61^and the log2 fold change was calculated. Statistical significance was determined by adjusted p-value <0.01. Gene Ontology and KEGG pathway enrichment analyses were performed based on the list of significant DGE genes.

### Statistical Analysis

Statistical analyses were performed using GraphPad Prism. Two-tailed Student’s t-test, One- or Two-way ANOVAs with post-hoc Tukey HSD test was performed. Statistical significance was denoted by * p-value <0.05, ** p-value <0.01, *** p-value <0.001, **** p-value <0.0001.

## References

1. Imbriano, C. & Belluti, S. Histone Marks-Dependent Effect on Alternative Splicing: New Perspectives for Targeted Splicing Modulation in Cancer? Int. J. Mol. Sci. 23, 8304 (2022).

2. Wang, Y. et al. Mechanism of alternative splicing and its regulation. Biomed. Rep. 3, 152–158 (2015).

3. Park, E., Pan, Z., Zhang, Z., Lin, L. & Xing, Y. The Expanding Landscape of Alternative Splicing Variation in Human Populations. Am. J. Hum. Genet. 102, 11–26 (2018).

4. Pan, Q., Shai, O., Lee, L. J., Frey, B. J. & Blencowe, B. J. Deep surveying of alternative splicing complexity in the human transcriptome by high-throughput sequencing. Nat. Genet. 40, 1413–1415 (2008).

5. Baralle, F. E. & Giudice, J. Alternative splicing as a regulator of development and tissue identity. Nat. Rev. Mol. Cell Biol. 18, 437–451 (2017).

6. Ke, S. & Chasin, L. A. Context-dependent splicing regulation. RNA Biol 8, 384–388 (2011).

7. Ganie, S. A. & Reddy, A. S. N. Stress-Induced Changes in Alternative Splicing Landscape in Rice: Functional Significance of Splice Isoforms in Stress Tolerance. Biology 10, 309 (2021).

8. Pagliarini, V., Naro, C. & Sette, C. Splicing Regulation: A Molecular Device to Enhance Cancer Cell Adaptation. BioMed Res. Int. 2015, 543067 (2015).

9. El Marabti, E. & Younis, I. The Cancer Spliceome: Reprograming of Alternative Splicing in Cancer. Front. Mol. Biosci. 5, 80 (2018).

10. Havens, M. A., Duelli, D. M. & Hastings, M. L. Targeting RNA splicing for disease therapy. Wiley Interdiscip. Rev. RNA 4, 247–266 (2013).

11. Ouyang, J. et al. The role of alternative splicing in human cancer progression. Am. J. Cancer Res. 11, 4642–4667 (2021).

12. Yeh, H.-S. & Yong, J. mTOR-coordinated Post-Transcriptional Gene Regulations: from Fundamental to Pathogenic Insights. J. Lipid Atheroscler. 9, 8–22 (2020).

13. Chang. J.-W., et al. mRNA 3’-UTR shortening is a molecular signature of mTORC1 activation. Nat Commun 6, 7218 (2015).

14. Cheng, S. et al. mTOR Contributes to the Proteome Diversity through Transcriptome-Wide Alternative Splicing. Int. J. Mol. Sci. 23, 12416 (2022).

15. Sun, J. et al. Dichotomous intronic polyadenylation profiles reveal multifaceted gene functions in the pan-cancer transcriptome. Exp. Mol. Med. 56, 2145–2161 (2024).

16. Szwed, A., Kim, E. & Jacinto, E. Regulation and metabolic functions of mTORC1 and mTORC2. Physiol. Rev. 101, 1371–1426 (2021).

17. Saxton, R. A. & Sabatini, D. M. mTOR Signaling in Growth, Metabolism, and Disease. Cell 168, 960–976 (2017).

18. Tian, T., Li, X. & Zhang, J. mTOR. Signaling in Cancer and mTOR Inhibitors in Solid Tumor Targeting Therapy. Int J Mol Sci 20, 755 (2019).

19. Gremke, N. et al. mTOR-mediated cancer drug resistance suppresses autophagy and generates a druggable metabolic vulnerability. Nat. Commun. 11, 4684 (2020).

20. Marcelo, K. L., Means, A. R. & York, B. The Ca(2+)/Calmodulin/CaMKK2 Axis: Nature’s Metabolic CaMshaft. Trends Endocrinol Metab 27, 706–718 (2016).

21. MacDonald, A. F. Concurrent regulation of LKB1 and CaMKK2 in the activation of AMPK in castrate-resistant prostate cancer by a well-defined polyherbal mixture with anticancer properties. BMC Complement Altern Med 18, 188 (2018).

22. Lin, F. et al. The camKK2/camKIV relay is an essential regulator of hepatic cancer. Hepatol. Baltim. Md 62, 505–520 (2015).

23. Juras, P. K. et al. Increased CaMKK2 expression is an adaptive response that maintains the fitness of tumor-infiltrating natural killer cells. Cancer Immunol. Res. 11, 109–122 (2023).

24. Fogarty, S. AMPK Causes Cell Cycle Arrest in LKB1-Deficient Cells via Activation of CAMKK2. Mol Cancer Res 14, 683–695 (2016).

25. Wang, Y. et al. Whole-exome sequencing combined with postoperative data identify c.1614dup (CAMKK2) as a novel candidate monogenic obesity variant. Front. Endocrinol. 15, 1334342 (2024).

26. Cao, W. Differential effects of PKA-controlled CaMKK2 variants on neuronal differentiation. RNA Biol 8, 1061–1072 (2011).

27. Hsu, L. S. Human Ca2+/calmodulin-dependent protein kinase kinase beta gene encodes multiple isoforms that display distinct kinase activity. J Biol Chem 276, 31113–31123 (2001).

28. Sabbir, M. G. CAMKK2-CAMK4 signaling regulates transferrin trafficking, turnover, and iron homeostasis. Cell Commun. Signal. CCS 18, 80 (2020).

29. Racioppi, L. & Means, A. R. Calcium/Calmodulin-dependent Protein Kinase Kinase 2: Roles in Signaling and Pathophysiology*. J. Biol. Chem. 287, 31658–31665 (2012).

30. Tokumitsu, H. & Sakagami, H. Molecular Mechanisms Underlying Ca2+/Calmodulin-Dependent Protein Kinase Kinase Signal Transduction. Int. J. Mol. Sci. 23, 11025 (2022).

31. Tokumitsu, H. & Sakagami, H. Molecular Mechanisms Underlying Ca2+/Calmodulin-Dependent Protein Kinase Kinase Signal Transduction. Int. J. Mol. Sci. 23, 11025 (2022).

32. Brzozowski, J. S. & Skelding, K. A. The Multi-Functional Calcium/Calmodulin Stimulated Protein Kinase (CaMK) Family: Emerging Targets for Anti-Cancer Therapeutic Intervention. Pharmaceuticals 12, 8 (2019).

33. Langendorf, C. G. et al. CaMKK2 is inactivated by cAMP-PKA signaling and 14-3-3 adaptor proteins. J. Biol. Chem. 295, 16239–16250 (2020).

34. Psenakova, K. et al. 14-3-3 protein directly interacts with the kinase domain of calcium/calmodulin-dependent protein kinase kinase (CaMKK2). Biochim. Biophys. Acta Gen. Subj. 1862, 1612–1625 (2018).

35. Ren, Y. & Shen, H.-M. Critical role of AMPK in redox regulation under glucose starvation. Redox Biol. 25, 101154 (2019).

36. Endo, H., Owada, S., Inagaki, Y., Shida, Y. & Tatemichi, M. Glucose starvation induces LKB1-AMPK-mediated MMP-9 expression in cancer cells. Sci Rep 8, 10122 (2018).

37. Liska, O. et al. TFLink: an integrated gateway to access transcription factor–target gene interactions for multiple species. Database 2022, baac083 (2022).

38. Miki, K. et al. Glucose starvation causes ferroptosis-mediated lysosomal dysfunction. iScience 27, 109735 (2024).

39. Li, L. et al. Ferroptosis is associated with oxygen-glucose deprivation/reoxygenation-induced Sertoli cell death. Int. J. Mol. Med. 41, 3051–3062 (2018).

40. Reichard, J. F., Motz, G. T. & Puga, A. Heme oxygenase-1 induction by NRF2 requires inactivation of the transcriptional repressor BACH1. Nucleic Acids Res. 35, 7074–7086 (2007).

41. Cai, L. et al. BACH1-Hemoxygenase-1 axis regulates cellular energetics and survival following sepsis. Free Radic. Biol. Med. 188, 134–145 (2022).

42. Reichard, J. F., Sartor, M. A. & Puga, A. BACH1 Is a Specific Repressor of HMOX1 That Is Inactivated by Arsenite. J. Biol. Chem. 283, 22363–22370 (2008).

43. Potteti, H. R. et al. Nrf2-AKT interactions regulate heme oxygenase 1 expression in kidney epithelia during hypoxia and hypoxia-reoxygenation. Am. J. Physiol.-Ren. Physiol. 311, F1025–F1034 (2016).

44. Li, R. et al. Glucose Starvation-Caused Oxidative Stress Induces Inflammation and Autophagy in Human Gingival Fibroblasts. Antioxid. Basel Switz. 11, 1907 (2022).

45. Endale, H. T., Tesfaye, W. & Mengstie, T. A. ROS induced lipid peroxidation and their role in ferroptosis. Front. Cell Dev. Biol. 11, (2023).

46. Panwar, V. Multifaceted role of mTOR (mammalian target of rapamycin) signaling pathway in human health and disease. Signal Transduct Target Ther 8, 375 (2023).

47. Chang, J.-W. et al. An Integrative Model for Alternative Polyadenylation, IntMAP, Delineates mTOR-modulated Endoplasmic Reticulum Stress Response - PubMed. Nucleic Acids Res. 46, 5996–6008 (2018).

48. Lee, G. et al. Post-transcriptional Regulation of De Novo Lipogenesis by mTORC1-S6K1-SRPK2 Signaling. Cell 171, 1545–1558.e18 (2017).

49. Passacantilli, I., Frisone, P., Paola, E., Fidaleo, M. & Paronetto, M. P. hnRNPM guides an alternative splicing program in response to inhibition of the PI3K/AKT/mTOR pathway in Ewing sarcoma cells. Nucleic Acids Res 45, 12270–12284 (2017).

50. Endo, H., Owada, S., Inagaki, Y., Shida, Y. & Tatemichi, M. Glucose starvation induces LKB1-AMPK-mediated MMP-9 expression in cancer cells. Sci. Rep. 8, 10122 (2018).

51. Bourouh, M. & Marignani, P. A. The Tumor Suppressor Kinase LKB1: Metabolic Nexus. Front. Cell Dev. Biol. 10, (2022).

52. Marcus, A. I. & Zhou, W. LKB1 regulated pathways in lung cancer invasion and metastasis. J. Thorac. Oncol. Off. Publ. Int. Assoc. Study Lung Cancer 5, 1883–1886 (2010).

53. Stockwell, B. R. et al. Ferroptosis: a regulated cell death nexus linking metabolism, redox biology, and disease. Cell 171, 273–285 (2017).

54. Yan, H. et al. Ferroptosis: mechanisms and links with diseases. Signal Transduct. Target. Ther. 6, 1–16 (2021).

55. Magtanong, L. et al. Context-dependent regulation of ferroptosis sensitivity. Cell Chem. Biol. 29, 1409–1418.e6 (2022).

56. Jang, N., Kim, I.-K., Jung, D., Chung, Y. & Kang, Y. P. Regulation of Ferroptosis in Cancer and Immune Cells. Immune Netw. 25, (2025).

57. Chen, X., Comish, P. B., Tang, D. & Kang, R. Characteristics and Biomarkers of Ferroptosis. Front. Cell Dev. Biol. 9, (2021).

58. Kim, D. et al. TopHat2: Accurate alignment of transcriptomes in the presence of insertions, deletions and gene fusions. Genome Biol. 14, (2013).

59. Bray, N. L., Pimentel, H., Melsted, P. & Pachter, L. Near-optimal probabilistic RNA-seq quantification. Nat. Biotechnol. 34, 525–527 (2016).

60. Karolchik, D., Hinrichs, A. S. & Kent, W. J. The UCSC Genome Browser. Curr Protoc Hum Genet Chapter 18, 18 6 1–18 6 33 (2011).

61. Varet, H., Brillet-Guéguen, L., Coppée, J.-Y. & Dillies, M.-A. SARTools: A DESeq2- and EdgeR-Based R Pipeline for Comprehensive Differential Analysis of RNA-Seq Data. PLoS One 11, 0157022 (2016).

